# Harnessing CRISPR-Cas9 for genome editing in *Streptococcus pneumoniae*

**DOI:** 10.1101/2020.06.13.149682

**Authors:** Dimitra Synefiaridou, Jan-Willem Veening

**Affiliations:** Department of Fundamental Microbiology, Faculty of Biology and Medicine, University of Lausanne, Biophore Building, CH-1015 Lausanne, Switzerland

**Author notes:** Correspondence to Jan-Willem Veening, Twitter: @JWVeening.

## Abstract

CRISPR systems provide bacteria and archaea with adaptive immunity against viruses and plasmids by detection and cleavage of invading foreign DNA. Modified versions of this system can be exploited as a biotechnological tool for precise genome editing at a targeted locus. Here, we developed a novel, replicative plasmid that carries the CRISPR-Cas9 system for RNA-programmable, genome editing by counterselection in the opportunistic human pathogen *Streptococcus pneumoniae*. Specifically, we demonstrate an approach for making targeted, marker-less gene knockouts and large genome deletions. After a precise double-stranded break (DSB) is introduced, the cells’ DNA repair mechanism of homology-directed repair (HDR) pathway is being exploited to select successful transformants. This is achieved through the transformation of a template DNA fragment that will recombine in the genome and eliminate recognition of the target of the Cas9 endonuclease. Next, the newly engineered strain, can be easily cured from the plasmid that is temperature-sensitive for replication, by growing it at the non-permissive temperature. This allows for consecutive rounds of genome editing. Using this system, we engineered a strain with three major virulence factors deleted. The here developed approaches should be readily transportable to other Gram-positive bacteria.

**Importance:** *Streptococcus pneumoniae* (the pneumococcus) is an important opportunistic human pathogen killing over a million people each year. Having the availability of a system capable of easy genome editing would significantly facilitate drug discovery and vaccine candidate efforts. Here, we introduced an easy to use system to perform multiple rounds of genome editing in the pneumococcus by putting the CRISPR-Cas9 system on a temperature-sensitive replicative plasmid. The here used approaches will advance genome editing projects in this important human pathogen.

## Introduction

*Streptococcus pneumoniae* (the pneumococcus) is a Gram-positive, human commensal that colonizes asymptomatically the mucosal surfaces of the upper respiratory tract (UTR) (Kadioglu et al. 2008). However, in susceptible groups like children, the elderly and the immunocompromised, it can occasionally become pathogenic causing diseases that range from a mild upper respiratory tract infection, acute otitis media and sinusitis, to a severe and potentially life-threatening condition such as pneumonia, bacteremia and meningitis (Simell et al. 2012). It is responsible for more than one million deaths annually (O’Brien et al. 2009) and in 2017, the World Health Organization (WHO) classified *S. pneumoniae* as one of twelve priority pathogens for which new antibiotics are urgently needed.

Historically, *S. pneumoniae* research played a central role in advancing molecular biology. While trying to develop a vaccine against the pneumococcus, Griffith discovered natural transformation (Griffith 1928). This was followed by research of Avery, MacLeod and McCarty to establish that DNA is the genetic material (Avery et al. 1944). Over the last decade, the pneumococcus has become a valuable model to study the cell biology of ovoid-shaped bacteria and several cell biological tools such as integration vectors, fluorescent reporters, inducible promoters and CRISPR interference have been established for this organism (Massidda et al. 2013; Keller et al. 2019; Liu et al. 2017). In addition, many selection and counterselection methods are available making it relatively easy to generate gene deletions, gene complementation mutants or point mutations in the pneumococcal genome (Sung et al. 2001; Halfmann et al. 2007; Y. Li et al. 2014; Sorg et al. 2019). However, all current gene deletion methods established for *S. pneumoniae* are poorly scalable and often require a specific genetic background to function (e.g. the *rpsL+* background in the janus system (Sung et al. 2001)).

In the case of gene replacement by selection markers, while powerful, this also has drawbacks, preventing further modifications of the genome when there are no further selectable markers available for additional strain development. Also, many important categories of gene mutation, such as missense substitutions and in-frame deletions, usually present no selectable phenotype (Sung et al. 2001). To circumvent these issues, we here established CRISPR genome editing for use as counterselection in the pneumococcus.

Clustered regularly interspaced short palindromic repeats (CRISPR) are present in many bacteria and most archaea (Jansen et al. 2002). Naturally, the system provides resistance against foreign genetic elements (e.g. phages or plasmids) via small noncoding RNAs that are derived from CRISPR loci. In class 2 type II CRISPR systems, the mature crRNA that is base-paired to a trans-activating crRNA (tracrRNA) forms a two-RNA structure that directs the CRISPR-associated proteins (e.g. Cas9 from *Streptococcus pyogenes*) to introduce a double-stranded break (DSB) into the target DNA locus. Site-specific cleavage occurs at locations determined by both base-pairing complementarity between the crRNA and the target protospacer DNA and a short protospacer adjacent motif (PAM) (Jinek et al. 2012). It has been demonstrated that the endonuclease can be programmed by engineering the mature dual-tracrRNA: crRNA as a single RNA chimera (sgRNA for single guide RNA), to cleave specific DNA sites. Thereby, modified versions of the system can be exploited as a biotechnological tool for precise, RNA-programmable genome targeting and editing (Jinek et al. 2012).

After the DSB has been introduced, the cell can utilize two major pathways in order to repair the break and survive: homologous recombination (HR) or non-homologous end-joining (NHEJ). In HR, a second intact copy of the broken chromosome segment, homologous to the DSB site, serves as a template for DNA synthesis across the break. In this mechanism, the crucial process of locating and recombining the homologous sequence is performed by RecA (Shuman and Glickman 2007). NHEJ does not rely on a homologous DNA template, as the two DNA ends are rejoined directly together. Most bacteria such as *S. pneumoniae* cannot perform NHEJ, while it is capable to perform HR (Prudhomme et al. 2002; 2014). DSB repair can be used as a way to generate mutants or desired changes to the genome by providing a HR template, and forms the basis of CRISPR engineering (Adli 2018). Indeed, early work, using integrative vectors and tracrRNAs, showed that Cas9 can be used to make markerless gene deletions in *S. pneumoniae* (Jiang et al. 2013).

In this study, we set out to establish a CRISPR engineering framework for *S. pneumoniae*. Specifically, we constructed a novel replicative plasmid containing a temperature-sensitive origin of replication (facilitating curing of the plasmid) carrying a genetic system for making targeted, marker-less gene knockouts and large genome deletions, which works with high efficiency in *S. pneumoniae*. The here developed plasmid system should be readily transportable to other Gram-positive bacteria as the used origin of replication was shown to be functional in *L. lactis* and *B. subtilis* (Bijlsma et al. 2007). While similar approaches have recently been undertaken to perform genome engineering in certain Gram-positive organisms such as *Enterococcus faecium* (de Maat et al. 2019), Clostridium (IC Cañadas et al. 2019) and *L. lactis* (Guo et al. 2019), a CRISPR-Cas9 gene editing system was not yet available for *S. pneumoniae* and the here described vector has the advantage of being readily curable due to its temperature sensitive origin of replication.

## Materials and Methods

### Bacterial strains, transformations and growth conditions

All pneumococcal strains used in this study are derivatives of the serotype 2 *S. pneumoniae* strain D39V (Avery et al. 1944, Slager et al. 2018). All plasmids where cloned in *NEB® Turbo Competent E. coli* (catalog number C2984; New England BioLabs). All the strains are shown in *Table 1*.

**Table 1:** Strain and plasmid list.

| *S. pneumoniae* Strains | Relevant Genotype | Reference |
| --- | --- | --- |
| D39V | Serotype 2 strain, wild type | Slager et al.,2018 |
| VL321 | *SPV_2146-P<sub>32</sub>-lacZ-chl-aliA* | This study |
| VL588 | *ssbB-luc-kan, cps::chl* | Lab collection |
| VL2172 | pPEPZ-read1-P<sub>3</sub>-BsaI-gfp-BsaI-read2-N701-p7 | Lab collection |
| VL3655 | D39V<br>+pDS05 [pG<sup>+</sup>host ori(Ts)-ermR-cloDF13ori-specR-P<sub>Zn_wtcas9</sub>-P<sub>3</sub>-gfp-sgRNA] | This study |
| VL3656 | *SPV_2146-lacZ-chl-aliA*<br>+pDS07 [pG<sup>+</sup>host ori(Ts)-ermR-cloDF13ori-specR-pZn_wtcas9-P<sub>3</sub>-sgRNA lacZ] | This study |
| VL3657 | $\Delta lacZ$<br>+pDS07 [pG<sup>+</sup>host ori(Ts)-ermR-cloDF13ori-specR-pZn_wtcas9-P<sub>3</sub>-sgRNA lacZ] | This study |
| VL3658 | $\Delta lacZ$ | This study |
| VL3659 | $\Delta cps$<br>+ pDS07 [pG<sup>+</sup>host ori(Ts)-ermR-cloDF13ori-specR-P<sub>Zn_wtcas9</sub>-P<sub>3</sub>-sgRNA lacZ] | This study |
| VL3660 | $\Delta cps$ | This study |
| VL3661 | $\Delta cps$<br>+ pDS12 [pG<sup>+</sup>host ori(Ts)-ermR-cloDF13ori-specR-P<sub>Zn_wtcas9</sub>-P<sub>3</sub>-sgRNA ply] | This study |
| VL3662 | $\Delta cps$, $\Delta ply$ + pDS12 [pG<sup>+</sup>host ori(Ts)-ermR-cloDF13ori-specR-P<sub>Zn_wtcas9</sub>-P<sub>3</sub>-sgRNA ply] | This study |
| VL3663 | $\Delta cps$, $\Delta ply$ | This study |
| VL3664 | $\Delta cps$, $\Delta ply$<br>+ pDS13 [pG<sup>+</sup>host ori(Ts)-ermR-cloDF13ori-specR-P<sub>Zn_wtcas9</sub>-P<sub>3</sub>-sgRNA lytA] | This study |
| VL3665 | $\Delta cps$, $\Delta ply$, $\Delta lytA$<br>+ pDS13 [$pG^+host$ $ori(Ts)$-$ermR$-$cloDF13ori$-$specR$-$P_{Zn\_wtcas9}$-$P_3\_sgRNA$ $lytA$] | This study |
| **Plasmids** |  |  |
| pDS05 | $pG^+host$ $ori(Ts)$-$ermR$-$cloDF13ori$-$specR$-$P_{Zn\_wtcas9}$-$P_3\_gfp$-$sgRNA$ | This study |
| pDS07 | $pG^+host$ $ori(Ts)$-$ermR$-$cloDF13ori$-$specR$-$P_{Zn\_wtcas9}$-$P_3\_sgRNA$ $lacZ$ | This study |
| pDS12 | $pG^+host$ $ori(Ts)$-$ermR$-$cloDF13ori$-$specR$-$P_{Zn\_wtcas9}$-$P_3\_sgRNA$ $ply$ | This study |
| pDS13 | $pG^+host$ $ori(Ts)$-$ermR$-$cloDF13ori$-$specR$-$P_{Zn\_wtcas9}$-$P_3\_sgRNA$ $lytA$ | This study |
| pRAS2 | pJWV01-$P_{32}$-$lacZ$ | Lab collection |

*S. pneumoniae* was grown either at 30°C, 37°C or 40°C (indicated) without shaking in liquid C+Y medium adapted from Adams and Roe (Martin et al. 1995) and contained the following compounds: adenosine (68.2 µM), uridine (74.6 µM), L-asparagine (302 µM), L-cysteine (84.6 µM), L-glutamine (137 µM), L-tryptophan (26.8 µM), casein hydrolysate (4.56 g L-1), BSA (729 mg L-1), biotin (2.24 µM), nicotinic acid (4.44 µM), pyridoxine (3.10 µM), calcium pantothenate (4.59 µM), thiamin (1.73 µM), riboflavin (0.678 µM), choline (43.7 µM), CaCl2 (103 µM), K2HPO4 (44.5 mM), MgCl2 (2.24 mM), FeSO4 (1.64 µM), CuSO4 (1.82 µM), ZnSO4 (1.58 µM), MnCl2 (1.29 µM), glucose (10.1 mM), sodium pyruvate (2.48 mM), saccharose (861 µM), sodium acetate (22.2 mM) and yeast extract (2.28 g L-1).

*E. coli* strains were cultivated in LB at 37°C with shaking. When appropriate, 100 μg/ml spectinomycin (spec) was added.

### Transformation

To transform the different plasmid variants into *S. pneumoniae*, cells were grown in C+Y medium (pH 6.8) at 37 °C to an OD_595_ of 0.1. Then, cells were treated for 12 min at 37°C with synthetic CSP-1 (100 ng mL^−1^) and incubated for 20 min at 30°C with the plasmid. After incubation, cells were grown in C+Y medium at the permissive temperature of 30°C for 120 min. *S. pneumoniae* transformants were selected by plating inside Columbia agar supplemented with 3% of defibrinated sheep blood (Thermo Scientific), followed by antibiotic selection, using erythromycin at concentration 0.25 μg/ml. Plates were incubated at 30°C.

To transform the HR template, cells were grown in C+Y medium (pH 6.8) at 30°C with 0.1 μg/ml erythromycin to an OD_595_ of 0.1. Then, cells were treated for 12 min at 37°C with synthetic CSP^−1^ (100 ng mL^−1^) and incubated for 20 min at 30°C with the HR template. After incubation, cells were grown in C+Y medium at 30°C for 20 min. Transformants were selected by plating inside Columbia agar supplemented with 3% of defibrinated sheep blood (Thermo Scientific), followed by CRISPR-mediated counter selection, using 1 mM ZnCl_2_/MnSO_4_. Plates were incubated at 30°C. Correct transformation was verified by PCR and sequencing. Working stocks of cells were prepared by growing cells in C+Y (pH 6.8), until an OD_595_ of 0.4. Cells were collected by centrifugation (1595 × *g* for 10 min) and resuspended in fresh C+Y medium with 15% glycerol and stored at −80°C.

### Plasmid curing

After the plasmid has been transformed into pneumococcus and successful deletion has been performed with the HR template and CRISPR-mediated counter selection, the plasmid can be eliminated from the pneumococcal cells. To achieve that, we first grow the strain with the plasmid at the non-permissive temperature, 40°C in C+Y, with a starting inoculum 1/10.000. Next, we plate the liquid culture in Columbia blood agar in several dilutions, to obtain single colonies after overnight incubation at 40°C. Single colonies were screened and 99% of them had successfully cured the plasmid from the strain.

### Recombinant DNA techniques

Oligonucleotides were ordered from Sigma and are listed in Table 2. Phanta Max Super-Fidelity DNA Polymerase (Vazyme) was used in PCR amplifications, restriction enzymes (ThermoFisher Scientific) were used for digestions and T4 DNA Ligase (Vazyme) was used for ligations.

**Table 2:**
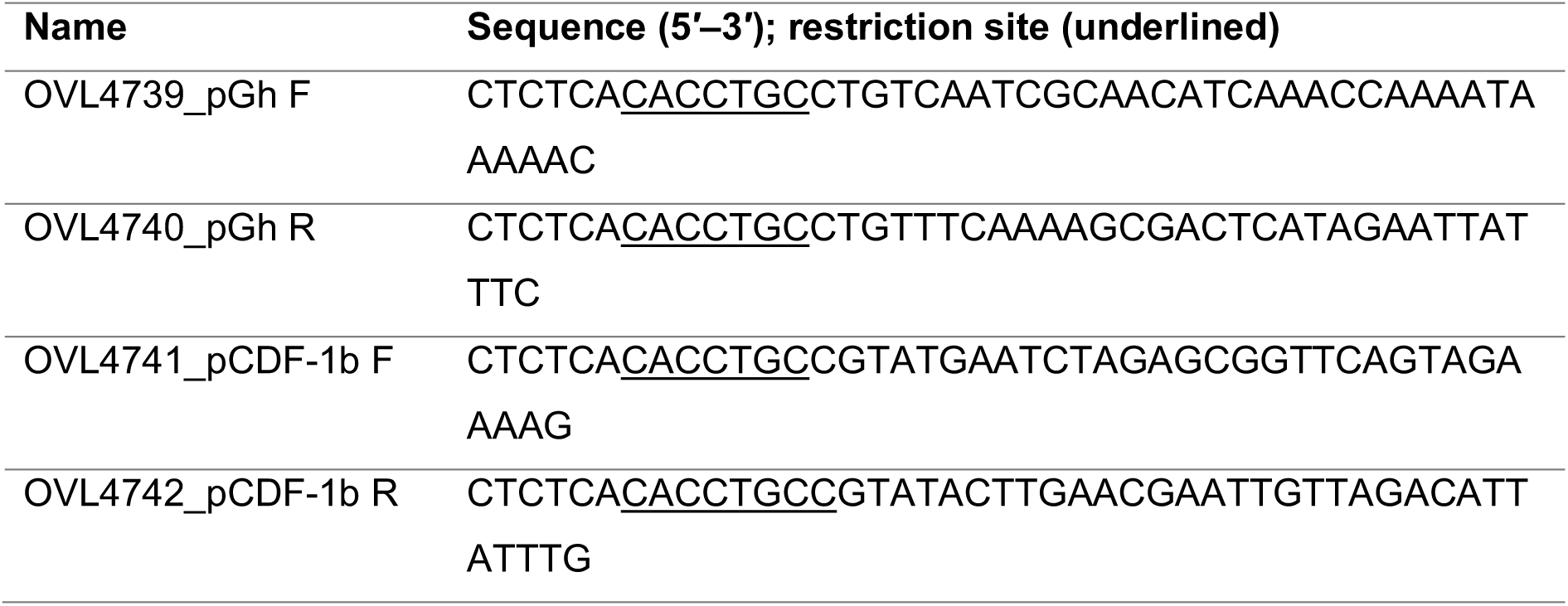
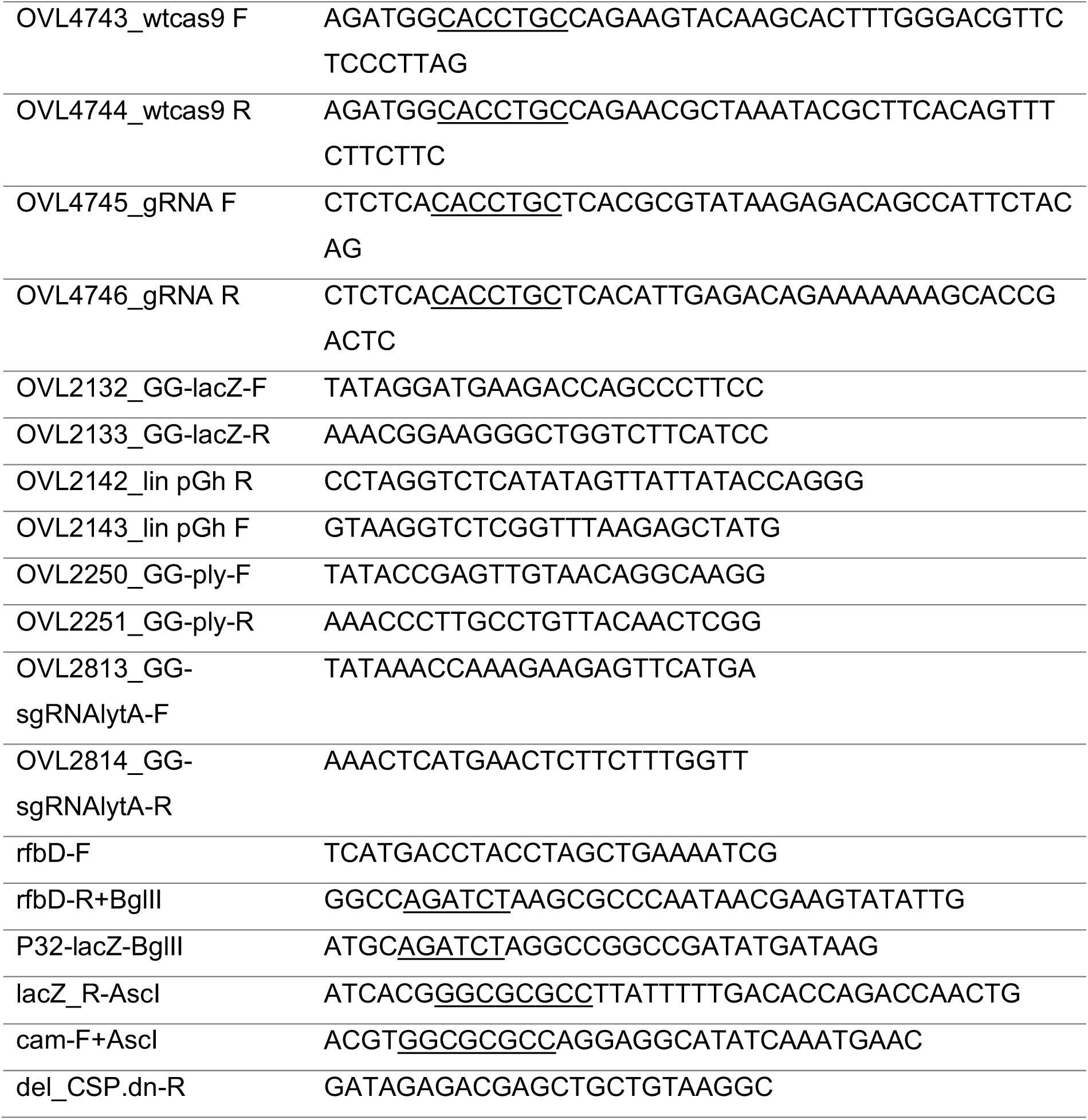
Oligonucleotides used in this study.

### Strain construction

pDS05 (*pG*^*+*^*host ori(Ts)-ermR-cloDF13ori-specR-P*_*Zn*_*_wtcas9-P*_*3*_*_gfp-sgRNA*). Gram-positive, temperature sensitive origin of replication *pG*^*+*^*host* (Maguin et al. 1992) and gene *ermR*, which confers resistance to erythromycin, were amplified from plasmid pGh9::ISS1 (Maguin et al. 1996) using the primers OVL4739_pGh F and OVL4740_pGh R (fragment 1). Gram-negative origin of replication *CloDF13* (CDF) and gene *specR*, which confers resistance to spectinomycin, were amplified from plasmid pCDF-1b (Nijkamp et al. 1986) with primers OVL4741_pCDF-1b F and OVL4742_pCDF-1b R (fragment 2). The gene which encodes wtCas9 under the control of the Zinc-inducible promoter was amplified from plasmid pJWV102-spCas9wt(van Raaphorst, Kjos, and Veening 2017), using the primers OVL4743_wtcas9 F and OVL4744_wtcas9 R (fragment 3). The sgRNA sequence in which the 20 base-pairing region of the sgRNA is replaced by the *gfp* gene was amplified from strain VL2172 with primers OVL4745_gRNA F and OVL4746_gRNA R (fragment 4). The four fragments were digested all together with restriction enzyme *AarI* and ligated. The ligation product was transformed into *E. coli* NEB Turbo and transformants were selected on LB agar with spectinomycin. Correct assembly was confirmed by PCR and sequencing.

pDS07 (*pG*^*+*^*host ori(Ts)-ermR-cloDF13ori-specR-P*_*Zn*_*_wtcas9-P*_*3_*_*sgRNA lacZ*). pDS05 was amplified with primers OVL2143_lin pGh F and OVL2142_lin pGh R (fragment 1). Spacer sequence of *sgRNA lacZ* was constructed by annealing primers OVL2132_GG-lacZ-F and OVL2133_GG-lacZ-R. Amplified pDS05 was digested with restriction enzyme *BsaI* and ligated with the annealed oligos. The ligation product was transformed into *E. coli* NEB Turbo and transformants were selected on LB agar with spectinomycin. Correct assembly was confirmed by PCR and sequencing.

pDS12 (*pG*^*+*^*host ori(Ts)-ermR-cloDF13ori-specR-P*_*Zn*_ *_wtcas9-P*_*3_*_*sgRNA ply*). pDS05 was amplified with primers OVL2143_lin pGh F and OVL2142_lin pGh R. Spacer sequence of *sgRNA ply* was constructed by annealing primers OVL2250_GG-ply-F and OVL2251_GG-ply-R. Amplified pDS05 was digested with restriction enzyme *BsaI* and ligated with the annealed oligos. The ligation product was transformed into *E. coli* NEB Turbo and transformants were selected on LB agar with spectinomycin. Correct assembly was confirmed by PCR and sequencing.

pDS13 (*pG*^*+*^*host ori(Ts)-ermR-cloDF13ori-specR-P*_*Zn*_ *_wtcas9-P*_*3_*_*sgRNA lytA*). pDS05 was amplified with primers OVL2143_lin pGh F and OVL2142_lin pGh R. Spacer sequence of *sgRNA lytA* was constructed by annealing primers OVL2813_GG-sgRNAlytA-F and OVL2814_GG-sgRNAlytA-R. Amplified pDS05 was digested with restriction enzyme *BsaI* and ligated with the annealed oligos. The ligation product was transformed into *E. coli* NEB Turbo and transformants were selected on LB agar with spectinomycin. Correct assembly was confirmed by PCR and sequencing.

VL321 (*SPV_2146*-P_32_-*lacZ-chl-aliA*). *rfbD* and *SPV_2146* were amplified from chromosomal DNA with primers rfbD-F and rfbD-R+BglII (fragment 1). P_32_-*lacZ* was amplified from pRAS2 (lab collection) with primers P32-lacZ-BglII and lacZ_R-AscI (fragment 2). A chloramphenicol resistance marker and *aliA* was amplified from strain VL588 with primers cam-F+AscI and del_CSP.dn-R. Fragment 1 was digested with restriction enzyme *Bgl*II, fragment 2 was digested with restriction enzymes *Bgl*II and *Asc*I and fragment 3 was digested with restriction enzyme *Asc*I. All three fragments were ligated together, and the ligation product was transformed into D39V. Correct assembly was confirmed by PCR and sequencing.

VL3655 (D39V + pDS05 [*pG*^*+*^*host ori(Ts)-ermR-cloDF13ori-specR-P*_*Zn*_*_wtcas9-P*_*3*_*_gfp-sgRNA*]). Plasmid pDS05 was transformed into D39V and transformants were selected on Columbia blood agar with erythromycin to produce the strain VL3655. The presence of the plasmid was confirmed by PCR and plasmid extraction.

VL3656 (*SPV_2146-lacZ-chl-aliA* + pDS07 [*pG*^*+*^*host ori(Ts)-ermR-cloDF13ori-specR-p*_*Zn*_*_wtcas9-P*_*3*_*_sgRNA lacZ*]). Plasmid pDS07, was transformed into VL321(*SPV_2146-lacZ-chl-aliA*) and transformants were selected on Columbia blood agar with erythromycin to produce the strain VL3656. The presence of the plasmid was confirmed by PCR and plasmid extraction.

VL3657 (Δ*lacZ* + pDS07 [*pG*^*+*^*host ori(Ts)-ermR-cloDF13ori-specR-P*_*Zn*_*_wtcas9-P*_*3*_*_sgRNA lacZ*]). HR template Δ*lacZ* was transformed into VL3656 and transformants were selected on Columbia blood agar with ZnCl_2_/MnSO_4_ to produce the strain VL3657. Correct integration was confirmed by PCR.

VL3658 (*ΔlacZ*). Strain VL3657 was cured from the plasmid, as described, resulting in strain VL3658.

VL3659 (Δ*cps* + pDS07 [*pG*^*+*^*host ori(Ts)-ermR-cloDF13ori-specR-P*_*Zn*_*_wtcas9-P*_*3*_*_sgRNA lacZ*]). HR template Δ*cps* was transformed into VL3656 and transformants were selected on Columbia blood agar with ZnCl_2_/MnSO_4_ to produce the strain VL3659. Correct integration was confirmed by PCR.

VL3660 (*Δcps*). Strain VL3659 was cured from the plasmid, as described, resulting in strain VL3660.

VL3661 (*Δcps* + pDS12 [*pG*^*+*^*host ori(Ts)-ermR-cloDF13ori-specR-P*_*Zn*_*_wtcas9-P*_*3*_*-sgRNA ply*]). Plasmid pDS12 was transformed into VL3660 and transformants were selected on Columbia blood agar with erythromycin to produce the strain VL3661. The presence of the plasmid was confirmed by PCR and plasmid extraction.

VL3662 (*Δcps, Δply* + pDS12 [*pG*^*+*^*host ori(Ts)-ermR-cloDF13ori-specR-P*_*Zn*_*_wtcas9-P*_*3*_*_sgRNA ply*]). HR template Δ*ply* was transformed into VL3661 and transformants were selected on Columbia blood agar with ZnCl_2_/MnSO_4_ to produce the strain VL3662. Correct integration was confirmed by PCR.

VL3663 (*Δcps, Δply*). Strain VL3662 was cured from the plasmid, as described, resulting in strain VL3663.

VL3664 (*Δcps, Δply* + pDS13 [*pG*^*+*^*host ori(Ts)-ermR-cloDF13ori-specR-P*_*Zn*_*_wtcas9-P*_*3*_*_sgRNA lytA*]). Plasmid pDS13 was transformed into VL3663 and transformants were selected on Columbia blood agar with erythromycin to produce the strain VL3664. The presence of the plasmid was confirmed by PCR and plasmid extraction.

VL3665 (*Δcps, Δply, ΔlytA* + pDS13 [*pG*^*+*^*host ori(Ts)-ermR-cloDF13ori-specR-P*_*Zn*_*_wtcas9-P*_*3*_*_sgRNA lytA*]). HR template Δ*lytA* was transformed into VL3664 and transformants were selected on Columbia blood agar with ZnCl_2_/MnSO_4_ to produce the strain VL3665. Correct integration was confirmed by PCR.

### Microscopy

*pneumoniae* cells were stored as exponential phase frozen cultures. Frozen stock was inoculated 1:100 in C+Y medium and pre-grown to OD600 ∼ 0.1. Cells were diluted once again 1:100 in fresh C+Y (with antibiotic, if applicable) and grown to exponential phase to achieve balanced growth.

Cells were grown as described above to achieve balanced growth and subsequently concentrated and brought onto a multi test slide carrying a thin layer of 1.2% agarose in C+Y. Imaging was performed on Fluorescence microscopy was performed on a Leica DMi8 through a 100x phase contrast objective (NA 1.40) with a SOLA Light Engine (lumencor) light source. Light was filtered through external excitation filters 470/40 nm (Chroma ET470/40x) for visualization of GFP. Light passed through a cube (Leica 11536022) with a GFP/RFP polychroic mirror (498/564 nm). External emission filters used were from Chroma ET520/40m. Images were captured using LasX software (Leica) and exported to ImageJ(Schneider, Rasband, and Eliceiri 2012) for final preparation.

Cell outlines were detected using MicrobeJ (Ducret et al. 2016). For all microscopy experiments, random image frames were used for analysis. The cell outline, object detection, and fluorescent intensity data were further processed using the R-package BactMAP (Raaphorst, Kjos, and Veening 2020).

### Transformation efficiency assays

To calculate the transformation efficiency, 1 μg/ml of PCR product of DNA fragment containing the upstream and the downstream region of the deletion target was added. Serial dilutions were plated either with or without 1 mM ZnCl_2_/MnSO_4_, and the transformation efficiency was calculated by dividing the number of transformants by the total viable count. All transformation efficiency values are averages of three biologically independent experiments.

### Whole genome sequencing and variant analysis

Genomic DNA was isolated using the FastPure Bacteria DNA Isolation Mini Kit (Vazyme) according to the manufacturers’ protocol and sent for whole genome sequencing. Illumina library prep and sequencing were carried out by Novogene (sequencing in PE150 mode). Reads were trimmed using Trimmomatic (Bolger, Lohse, and Usadel 2014), then assembled using SPAdes (Nurk et al. 2013) and contigs were reordered in Mauve (Darling et al. 2004) using D39V as a reference (Slager et al NAR 2018). Reads were mapped onto the scaffold using bwa (H. Li and Durbin 2009), read depth was determined in samtools (H. Li et al. 2009), and plotted in R (Team 2014). In order to detect small variants, raw reads were mapped onto the reference using bwa, and variants were detected using Freebayes (Garrison and Marth 2012). Potential variants with PHRED scores greater than 30 were filtered out on DP >5 using vcflib (Garrison n.d.), then intersected with the D39V annotation using Bedtools (Quinlan and Hall 2010)

## Results

### A novel *S. pneumoniae* replicative plasmid that carries the CRISPR-Cas9 system

First, a replicative plasmid was designed and constructed (Figure 1), by combining PCR amplified genomic and plasmid parts. The main idea behind the choice for individual vector components relied in creating a platform with the CRISPR-Cas9 system in *S. pneumoniae* while at the same time allowing for plasmid propagation in both Gram-positive and Gram-negative hosts. The modular vector consists of six individual components. Two origins of replication; the high-copy pG^+^host replicon, which is a replication thermosensitive derivative of pWV01 (Otto et al. 1982) that in *L. lactis*, (and other Gram-positive bacteria) replicates at 28^°C^ but is lost above 37^°C^, and the low-copy *CloDF13* (CDF) replicon for propagation in *E. coli*. By combining these two origins of replication, it ensures low copy numbers at 37^°C^ in *E. coli* thereby preventing toxicity of the CRISPR-Cas9 system while cloning. Additionally, it has the gene which encodes wtCas9 under the control of the Zinc-inducible promoter Pzn (Eberhardt et al. 2009) (plasmid pDS05) and genes conferring spectinomycin (*E. coli*) and erythromycin (*S. pneumoniae*) resistance. Finally, it has the strong synthetic constitutive P3 promoter (Sorg et al. 2015) driving the sgRNA sequence in which the 20 base-pairing region of the sgRNA is replaced by the *gfp* gene flanked by two *Bsa*I restriction sites. This allows for easy replacement of *gfp* by the spacer sequence of sgRNA with golden gate cloning. This way, successful cloning of the sgRNA allows for easy selection by absence of GFP fluorescence, giving us a versatile vector for different sgRNAs (see below).

**Figure 1:**
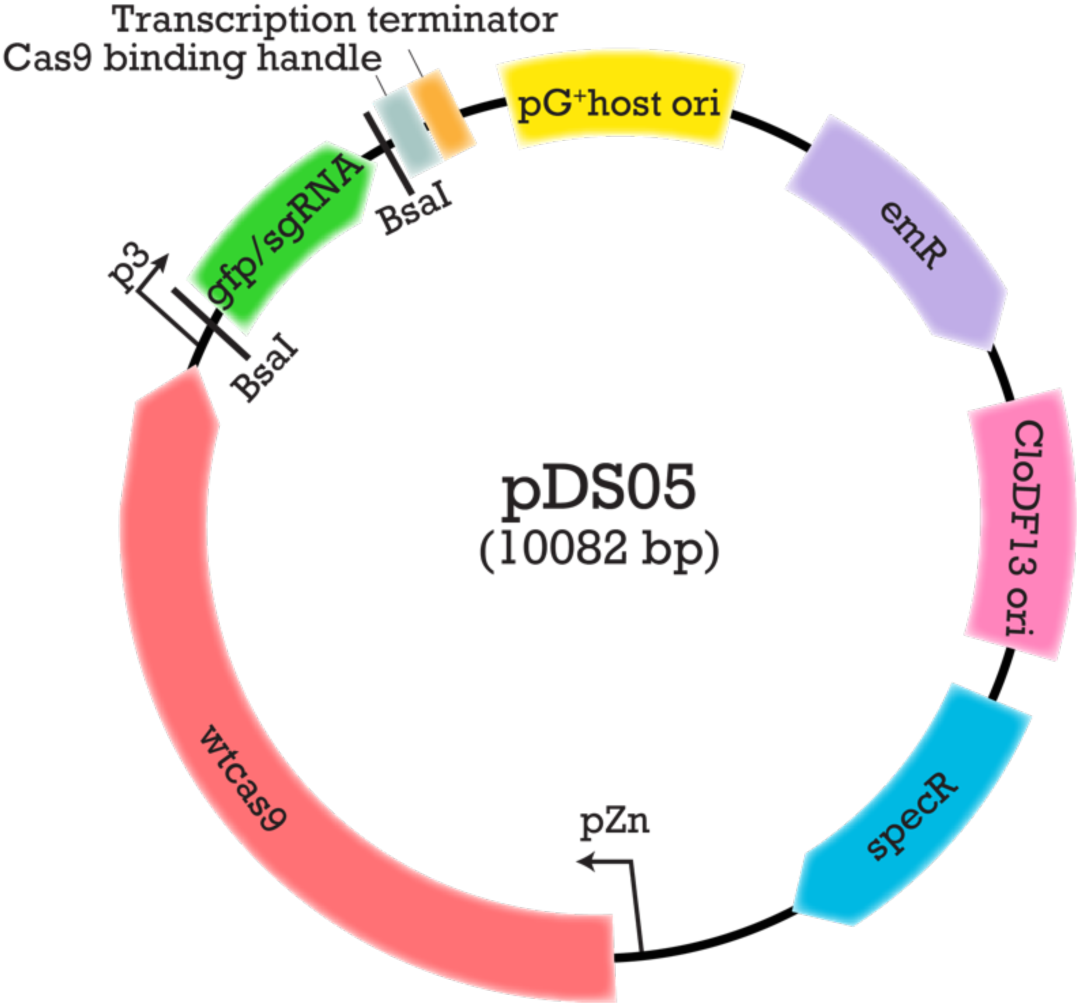
Schematic representation of plasmid pDS05.

### Successful plasmid propagation and plasmid curing in *S. pneumoniae*

To test that the newly constructed plasmid was being replicated and genes were expressed in *S. pneumoniae*, we grew strain VL3655 (carrying plasmid pDS05, see Fig. 1), which encodes GFP, in C+Y medium at 28°C in the presence of erythromycin. Fluorescence microscopy demonstrated that all cells produced GFP, although significant cell-to-cell variability was observed (Figure 2a). GFP intensity levels were determined in exponentially growing cells (balanced growth). Additionally, cells pre-grown in 28°C in presence of erythromycin (T=0) were split and grown under four different conditions. The permissive 28°C with and without erythromycin in the growth medium and the non-permissive 40°C with and without erythromycin. Note that growth was balanced by re-diluting exponentially growing cells several times. Cells were being collected every two hours for 8 hours and GFP intensity levels were determined using fluorescence microscopy and images were analyzed using MicrobeJ and BactMAP (Ducret et al. 2016; Raaphorst et al. 2020) (Figure 2b and c). The results show that GFP levels and, by extension plasmid copy number, stay stable at 28°C with erythromycin, and slowly decreases in the absence of antibiotic pressure. Furthermore, GFP levels decrease significantly in cells grown at 40°C, confirming that this is a non-permissive temperature for propagation of the plasmid. Absence of antibiotic pressure seems also to facilitate the decrease of the intensity levels of GFP, suggesting that the plasmid gets eliminated successfully under these conditions.

**Figure 2:**
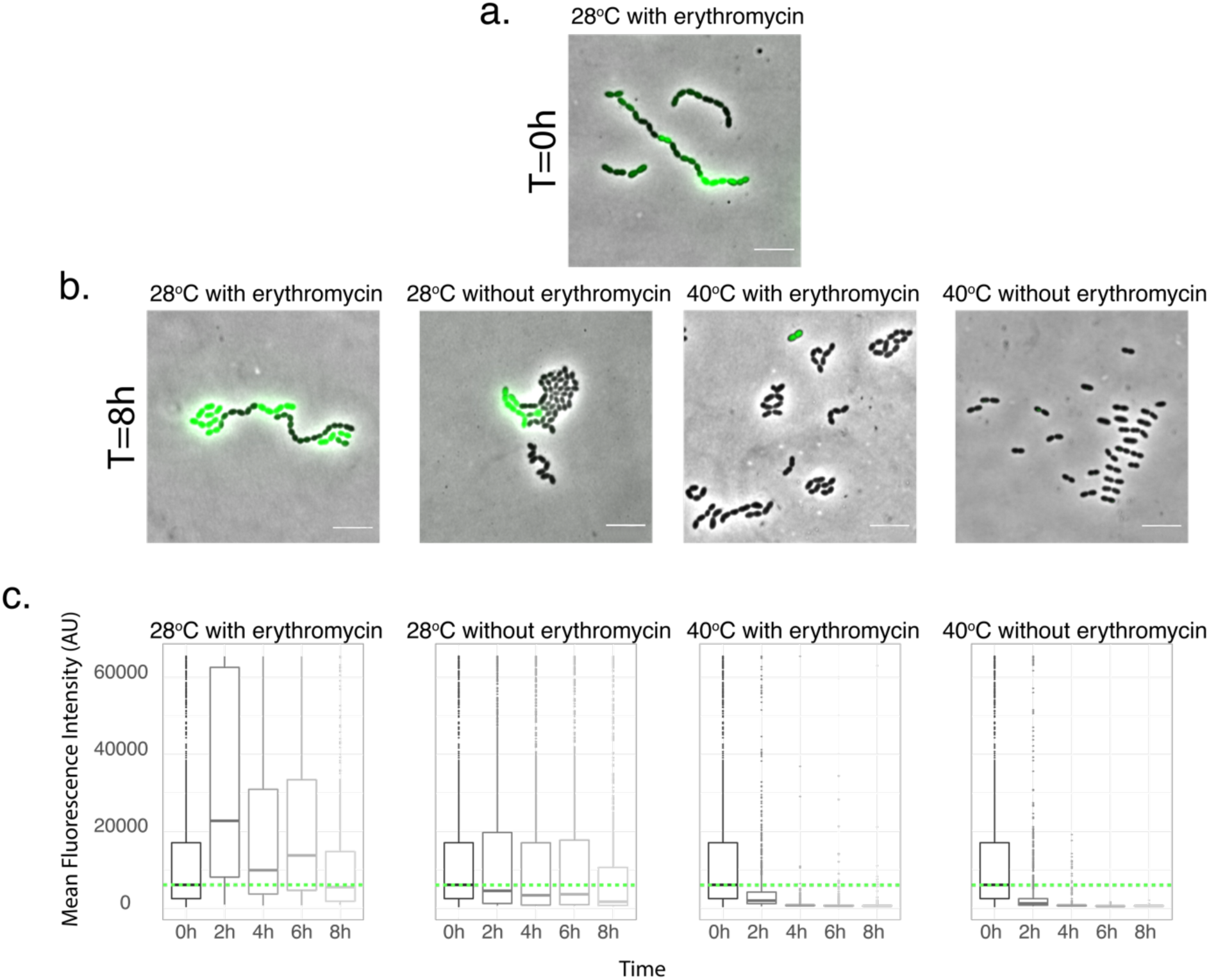
Microscopy analysis of strain VL3655 (D39V, pDS05). Overlay of GFP signals with phase contrast image shows GFP expression. a. Preculture grown at 28°C with erythromycin (T=0h), b. Images are shown of cells grown for 8h as exponentially growing cells (balanced growth) in four different conditions: 28°C with erythromycin, 28°C without erythromycin, 40°C with erythromycin, and 40°C without erythromycin. Scale bar in all images = 6 μm. c. Quantification of mean fluorescence intensity of GFP of cells grown under four different conditions: 28°C with erythromycin, 28°C without erythromycin, 40°C with erythromycin, and 40°C without erythromycin) in time points 0, 2, 4, 6 and 8 hours after dilution from the 28°C with erythromycin condition. Fluorescence microscopy of ±1000 cells per condition per time point were quantified and analyzed using MicrobeJ and BactMap and plotted as box plots, (box size and line represent the average intensity per cell) (see Materials and Methods). The green dotted horizontal line indicates the mean fluorescence of cells from the preculture harboring pDS05.

## CRISPR/Cas9-Mediated Counterselection

Once a deletion target has been selected, the plasmid with the specific sgRNA needs to be constructed. The targeting of Cas9 to a locus of interest is achieved by cloning two annealed 24-bp DNA oligonucleotides (containing the 20 bp protospacer element) into the sgRNA backbone that matches the specified locus. First, *gfp* is removed from pDS05 by digesting with *Bsa*I. Complementary oligos that carry the spacer sequence are annealed together. They are designed in a way that after annealing, they have overhangs complementary with those left after digestion of the backbone, as previously described (Liu et al. 2020) (Figure 3). The desired plasmid is obtained after ligation and transformed to *E. coli*. False positive transformants are easily identified, since they still carry the *gfp* and produce detectable fluorescent green colonies.

**Figure 3:**
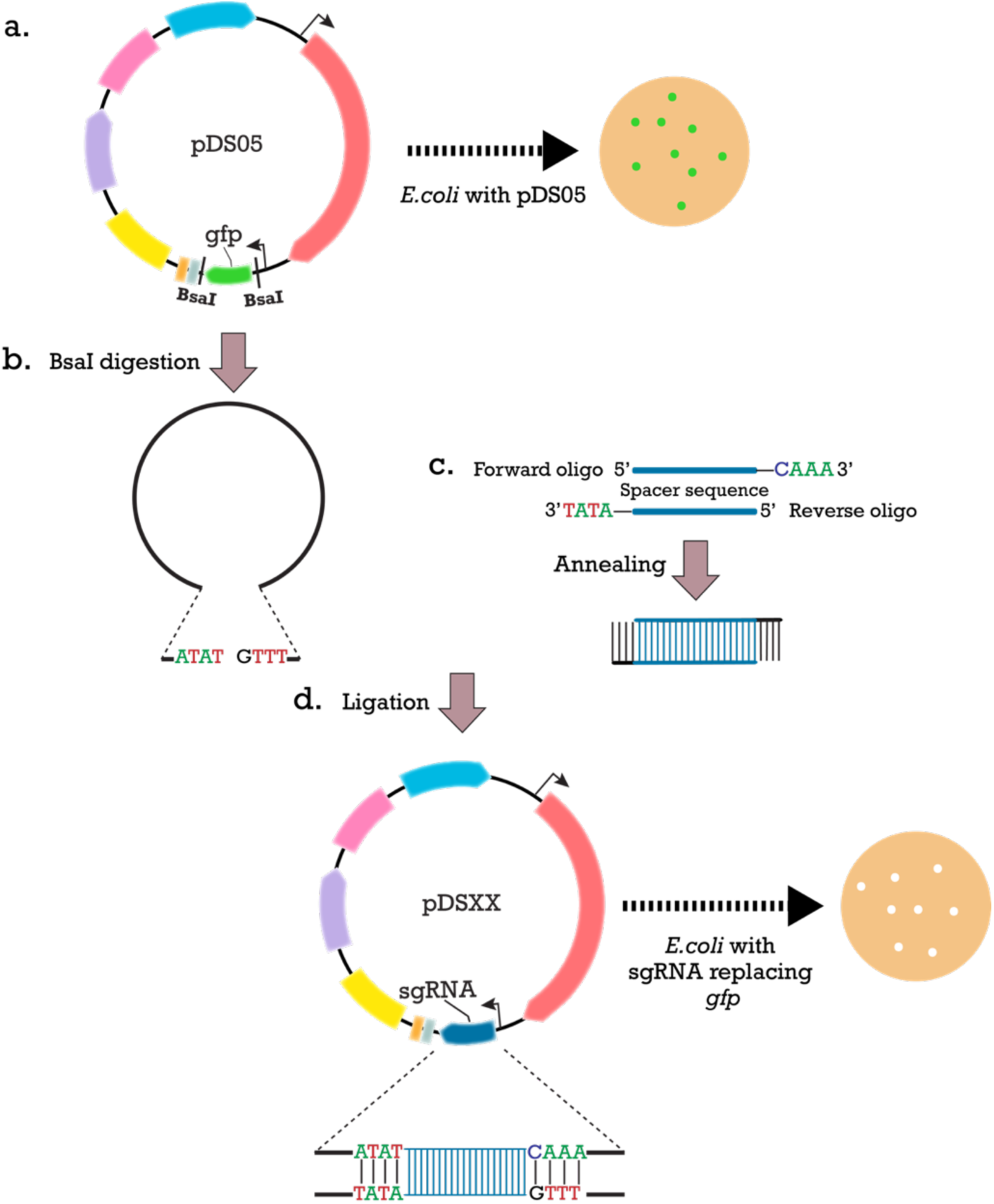
Workflow of sgRNA replacement. a. pDS05 was designed to facilitate easy replacement of *gfp* by the spacer sequence of the desired sgRNA with golden gate cloning, allowing also for detection of false positive transformants. *gfp*, which encodes a green fluorescent protein, is in place of the spacer sequence of sgRNA and flanked by *Bsa*I sites. *E. coli* with pDS05 produces green fluorescent colonies b. *Bsa*I digestion of the vector exposes 4 nt overhangs c. For each sgRNA, forward and reverse oligos were designed, as a reverse complement of each other, which after being annealed together, were containing the 20 bp spacer sequence and 4 nt overhangs, that can be specifically annealed with the digested vector. d. Ligation of the digested vector with the sgRNA annealed product was transformed into *E. coli*, producing white colonies.

After isolating the plasmid from *E. coli*, it needs to enter the pneumococcal cells. To achieve this, competence is induced by adding synthetic CSP and the transformation machinery is utilized. Note that competence-dependent transformation with a replicative plasmid is less efficient than transforming with linear homologous DNA (Johnston et al. 2014), so transformation efficiencies with pDS05 are typically low. However, in this case, only 1 successful transformant is required. Next, an HR template is constructed that consists of the upstream and downstream region of the deletion target. Following again induction of competence by CSP, the pneumococcal cells harboring the pDS05-derivative are transformed with this template. Transformants are selected by plating with ZnCl_2_/MnSO_4_, the inducer of Cas9, offering CRISPR-mediated counter-selection. Only cells that have taken up and integrated the HR template, thereby eliminated the recognition target of the sgRNA/Cas9 complex, would be able to survive, while untransformed cells will undergo DNA cleavage and die (Figure 4).

**Figure 4:**
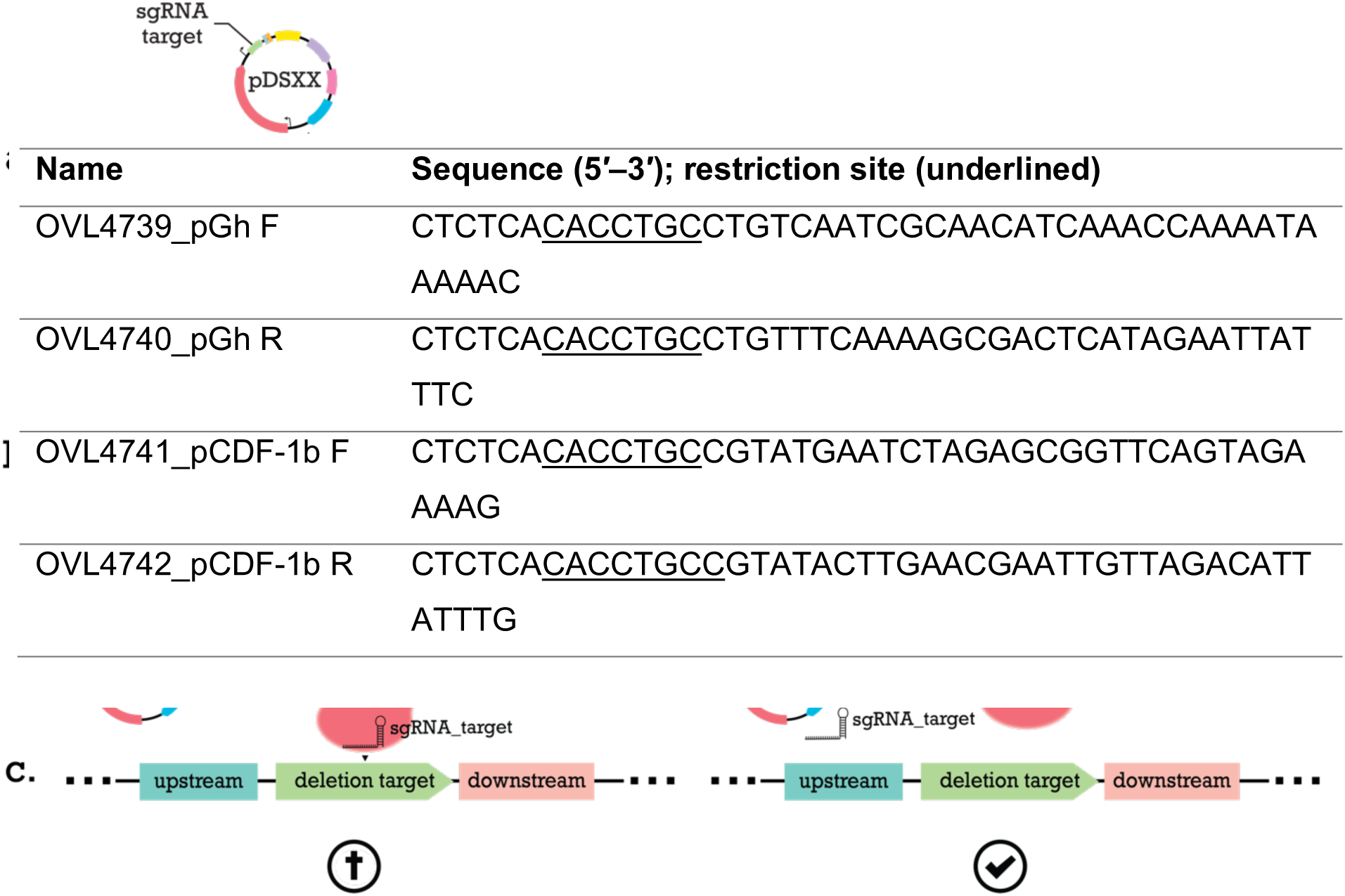
Workflow for markerless deletions. a. Uptake of the plasmid by the strain with the sgRNA sequence for the desired deletion. b. Transformation of a homology recombination (HR) template consisting of the ligation of the upstream and downstream region of the deletion target. c. Plating transformants with Zn^2+^ to induce expression of Cas9. Only the cells that have taken up the HR template eliminating the recognition target are able to survive.

### Deleting genes and large chromosomal regions from the *S. pneumoniae* genome

To assess the efficiency of the system, we first constructed a strain (strain VL3656; Figure 5a) in which we placed the *E. coli lacZ* gene under a constitutive promoter behind the *S. pneumoniae* D39V *cps* locus (encoding the capsule). *lacZ* encodes for a β-galactosidase that hydrolyzes X-gal to produce a blue product, allowing for blue/white screening on plates. Colonies with blue color would still carry the *lacZ* gene, while colonies with the standard white/green (on blood agar) color would indicate that the gene has been deleted from the chromosome.

**Figure 5:**
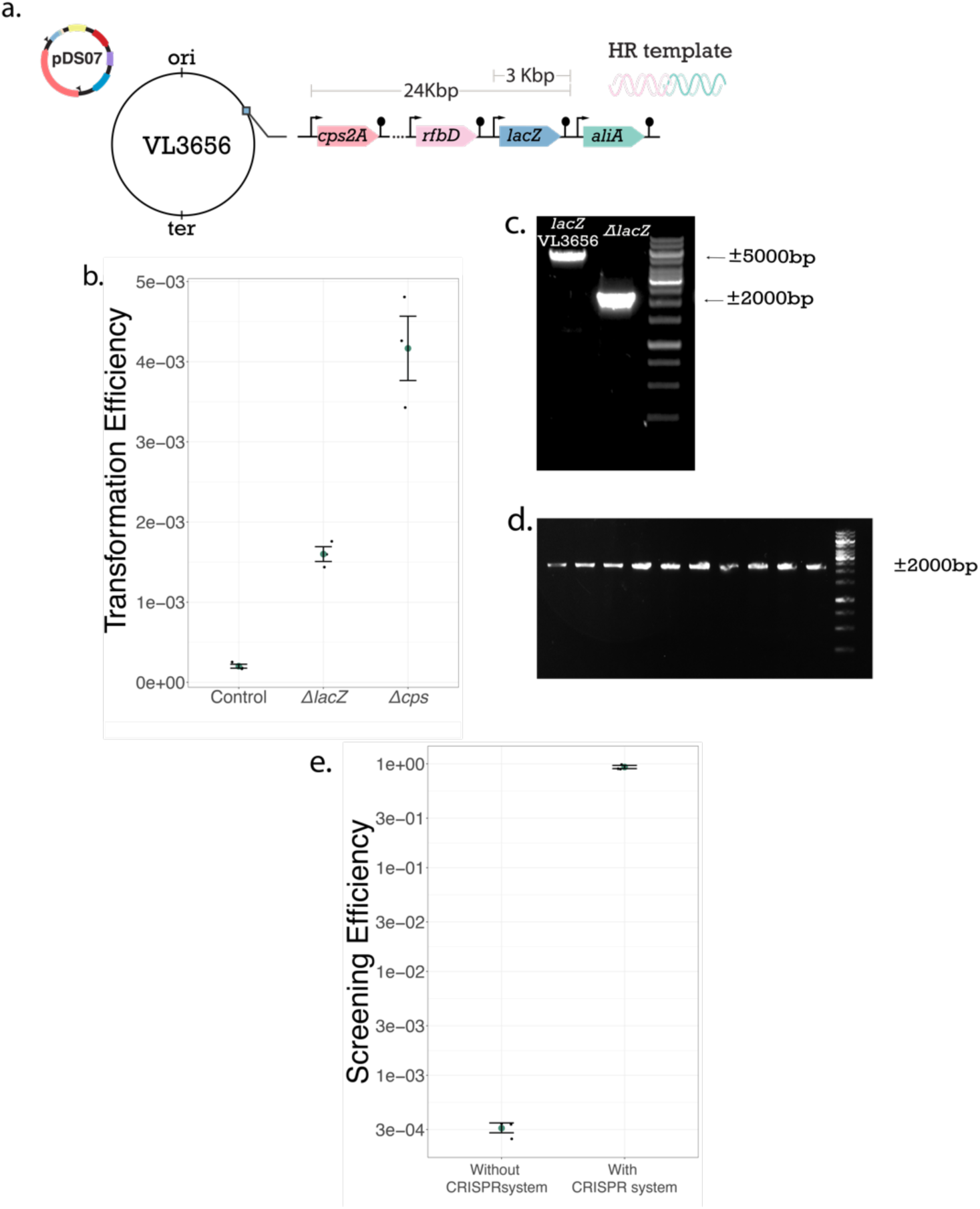
Genome editing in *S. pneumoniae* using CRISPR/Cas9. a. Schematic representation of strain VL3656. The *lacZ* gene has been inserted downstream the capsule operon and a version of the plasmid with a sgRNA targeting *lacZ* has been transformed in the strain. Control is the transformation assay of strain VL3656 in the absence of HR template DNA. b. Transformation efficiency of *lacZ* and capsule operon deletion. The transformation efficiency was calculated by dividing the total number of cells as counted on plates without Cas9 inducer (1 mM Zn^2+^) by the number of colonies in the presence of inducer. c. Colony PCR analysis of expected sizes. d. Eight randomly selected transformants of *lacZ* deletion. e. Efficiency of successful transformants screened for integration of the *lacZ* deletion when using no selection and when using the CRISPR system. Data represent the average of three independent experiments (± SE).

Strain VL3656 also carries the pDS07 plasmid, which contains a sgRNA targeting *lacZ*. Next, we constructed an HR template that consisted of the 1000 bp upstream and 1000 bp downstream region of *lacZ* (excluding *lacZ*) (**Figure *5***a) and we transformed VL3656 with this template. Transformants were selected by plating on agar containing ZnCl_2_ to induce expression of Cas9.

After transformation of strain VL3656 with the HR template, transformation efficiency was calculated (Figure 5b). The CRISPR-mediated counter-selection, offered by the system, worked successfully. The selection efficiency was high and almost all colonies in the transformation where the HR template was given and Cas9 was induced had their original color, indicating the *lacZ* gene has been successfully deleted.

Transformants were tested for correct deletion of *lacZ* by colony PCR. The primers used were binding 1000bp upstream and downstream of *lacZ*, setting the correct PCR product of the successful deletion at the 2000bp (5000bp if *lacZ* was not deleted) (Figure 5c). All the tested colonies had the expected product demonstrating successful deletion of *lacZ*, resulting in strain VL3657 (Figure 5d).

Additionally, we also used the system to delete an even larger chromosomal fragment. For this, we targeted the operon that encodes the capsule and the *lacZ* gene that had been inserted downstream of it, which is around 24Kbp long, allowing for blue/white screening. Once again, selection efficiency was very high and almost all colonies had their original color (Figure 5b). Colony PCR verified correct deletion of the *cps-lacZ* chromosomal region (see below).

Using the same HR template to delete *lacZ*, we also performed transformation assays without the counterselection offered, by inducing our CRISPR system (Figure 5f). Thousands of colonies needed to be screened to find successful transformants with the original colony color, among the blue colonies. In contrast, by using the system, almost with absolute success rate, all the colonies on our plates are the correct transformants, demonstrating how efficient our system is to easily select edited cells.

### Consecutive deletions of virulence factors of *S. pneumoniae*

Once the capsule operon and *lacZ* were removed from the chromosome, it was confirmed by colony PCR. All tested colonies demonstrated the expected PCR product. One such colony was picked resulting in strain VL3659. Next, we grew the new strain at the non-permissive temperature (40°C), eliminating the plasmid, resulting in strain VL3660 (*Δcps*).

To examine whether the system could be used in multiple rounds of genome editing, we attempted to delete the virulence factor pneumolysin. To delete the *ply* gene, we designed a sgRNA targeting pneumolysin and constructed plasmid pDS12, which we transformed into VL3660. Following the same procedure as used to delete the *cps* operon, we deleted *ply*. Again, to confirm the successful deletion, the same principle for the primers set was used. All the colonies from the transformation plate had the expected PCR product demonstrating extremely high selection efficiency using the CRISPR-Cas9 system. Finally, following the same strategy, we also deleted another important virulence factor, *lytA* resulting in strain VL3665 (**Figure *6***d; *Δcps, Δply, ΔlytA +* pDS13).

### Cas9-dependent genome editing is specific without evidence for off-target cutting in *S. pneumoniae*

After three consecutive deletions, using our novel plasmid with the CRISPR-Cas9 system, the final result was strain VL3665. It has been previously shown that Cas9 tolerates mismatches between guide RNA and target DNA at different positions in a sequence-dependent manner, resulting in off-target DSB (Hsu et al. 2013). To examine the fidelity of our CRISPR system and whether there were detectable genome-wide off-target effects, we performed whole-genome sequencing (WGS). The analysis detected one single SNP in the genome, in the gene *psaA* (*SPV_1463*), a manganese ABC transporter. The mutation results in a D137E amino acid change. Using Sanger sequencing we confirmed that this SNP occurred only in the last strain of the consecutive deletions and it has not been present in the intermediate steps. There is no evidence to believe that this mutation is associated with an off-target effect as the sequence surrounding the SNP is completely different from the used sgRNA present in plasmid pDS13 and most probably happened randomly during growth, without affecting the fitness (**Error! Reference source not found**.).

On strain VL3665, with the three major virulence factors deleted, we performed the final confirmations. Colony PCR showed that the chromosomal fragments have successfully been deleted from the chromosome (Figure 6a and b). Additionally, reads from whole-genome sequencing were competitively mapped onto the reference genome, our wild type lab strain D39V (Figure 6c). Direct comparison between the genomes reveals the three chromosomal positions that the deletions have taken place, since in these positions, the chromosomal dosage drops. Therefore, we confirmed that we had successfully performed markerless deletions of these three genes (Figure 6d).

**Figure 6:**
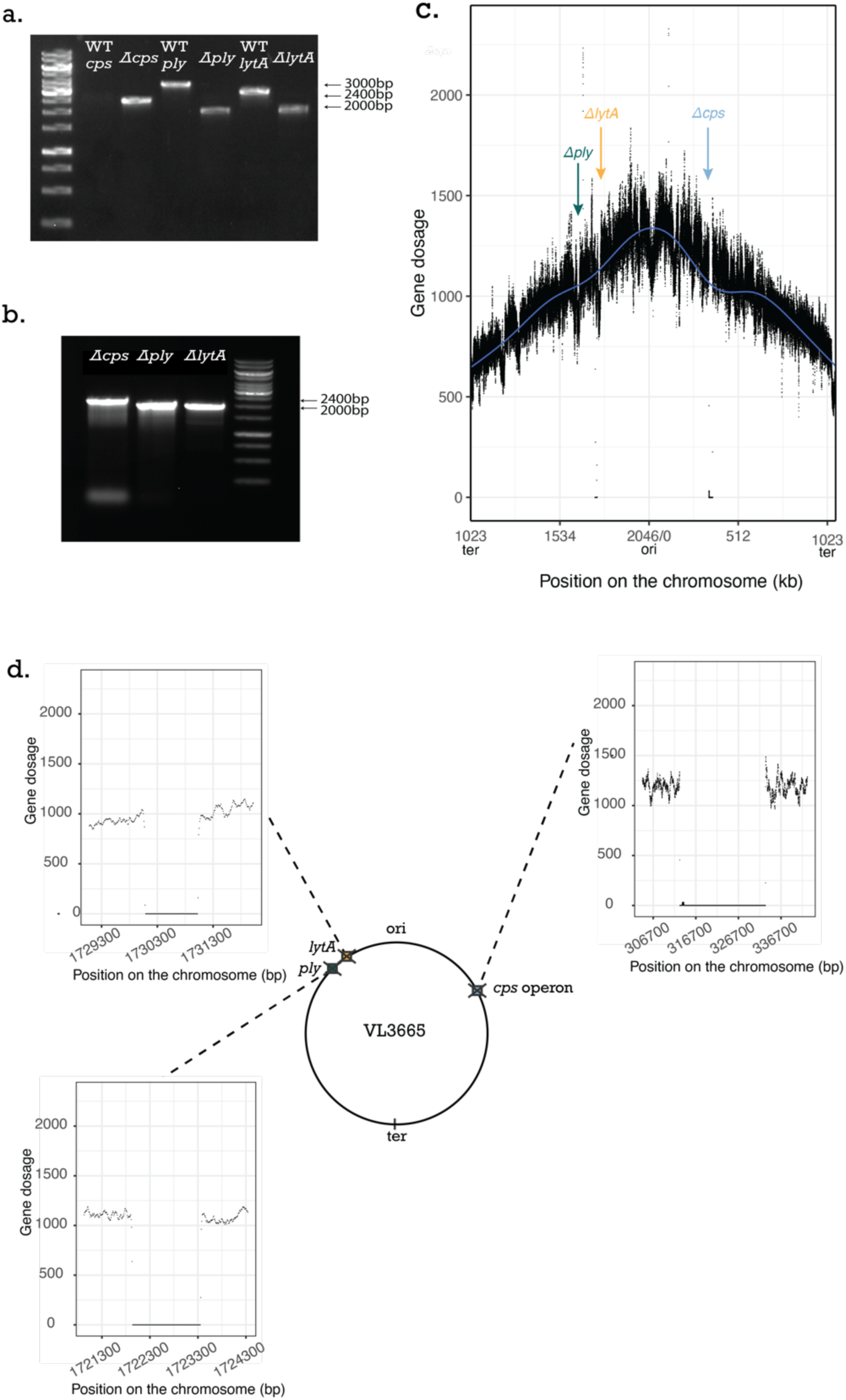
Genome analysis of the Δ*cps*, Δ*ply*, Δ*lytA* triple mutant generated using CRISPR-Cas9 editing. a. Colony PCR analysis of expected sizes for deletion of three virulence genes. WT vs VL3665. b. Colony PCR analysis of three virulence gene deletions in the final strain VL3665. c. Whole genome marker frequency analysis of strain VL3665. d. Schematic representation of strain VL3665 with three virulence gene deletions and zoom in 10kb upstream and downstream of the regions we deleted. The number of mapped reads (gene dosage) is plotted as a function of the position on the circular chromosome.

## Discussion

Genetic manipulation of microorganisms has been pivotal for the development of biotechnological tools and the study of microorganisms themselves. In this study, we have developed a novel, replicative plasmid with a temperature-sensitive origin of replication carrying a CRISPR-Cas9 based system for advanced and markerless genome engineering in the bacterium *S. pneumoniae*. In particular, we demonstrate that we have successfully deleted genes and large chromosomal regions in a precise and sequential way.

The here designed plasmid has the temperature sensitive origin of replication *pG*^*+*^*host*, which is a derivative of pWV01 of *L. lactis* and can be successfully propagated in pneumococcus at 30°C, while it is not stable at 40°C. Indeed, we show that our *pG*^*+*^*host* derivative, pDS05, is rapidly lost at 40°C (Fig. 2). We used this feature to eliminate the plasmid from the strains, upon the desired deletion. The fact that the copies of the plasmid vary per cell does not affect our system, since even one copy of *cas9* seems to be sufficient to perform the DSB (van Raaphorst et al. 2017).

Specifically, our approach is to harness this CRISPR and the homologous recombination system, to perform CRISPR/Cas9-Mediated Counterselection. Following the same principle of transformation with antibiotic selection, successful transformants survive the CRISPR/Cas9 induced DSB, like they survive growth in antibiotics, if they uptake the rescue HR template. Applying this, our CRISPR system manages to select for transformants in which single genes or even large chromosomal regions were deleted with very high efficiency. Comparing this to just performing natural transformation without any counterselection, which would be an alternative for clean deletions, we show the advantages of our system (Fig. 4f). Without it, we would need to screen many colonies to find correct transformants, depending on the target. This will have to be done by colony PCR, since in most of the cases, the desired deletion will not give any phenotypic difference in the colonies of the successful transformant, which is a costly and time demanding process. On the other hand, with the CRISPR/Cas9-Mediated counterselection, nearly all the colonies that we obtained were the desired transformant, since very few false positives have been observed.

Since we are ultimately interested to remove multiple genes and chromosomal regions from the genome, we also needed to demonstrate that our system is capable of consecutive deletions. The key for this was to easily eliminate the plasmid from the newly constructed strain. By growing the strain still carrying the plasmid at the high, non-permissive temperature, we manage to easily cure it. Next, we can transform a new plasmid and proceed further with our deletions. Specifically, after we deleted the capsule, we next deleted virulence factors *ply* and *lytA*, proving that our CRISPR-Cas9 system has flexibility in genetic manipulation of the bacterial genome. Together, the here described plasmid and approach will be useful for the pneumococcal research community and may be applicable to other Gram-positive bacteria as well. Plasmid pDS05 is available from Addgene (accession number pending).

## Acknowledgements

We are grateful to Paddy Gibson for help with WGS, Renske van Raaphorst for help with image analysis and all the members of the Veening laboratory for stimulating discussions. Work in the Veening lab is supported by the Swiss National Science Foundation (SNSF) (project grant 31003A_172861), a JPIAMR grant (40AR40_185533) from SNSF and ERC consolidator grant 771534-PneumoCaTChER.

