## Supplementary Figure S1 for "Harnessing CRISPR-Cas9 for genome editing in *Streptococcus pneumoniae*"

### Supplementary figures

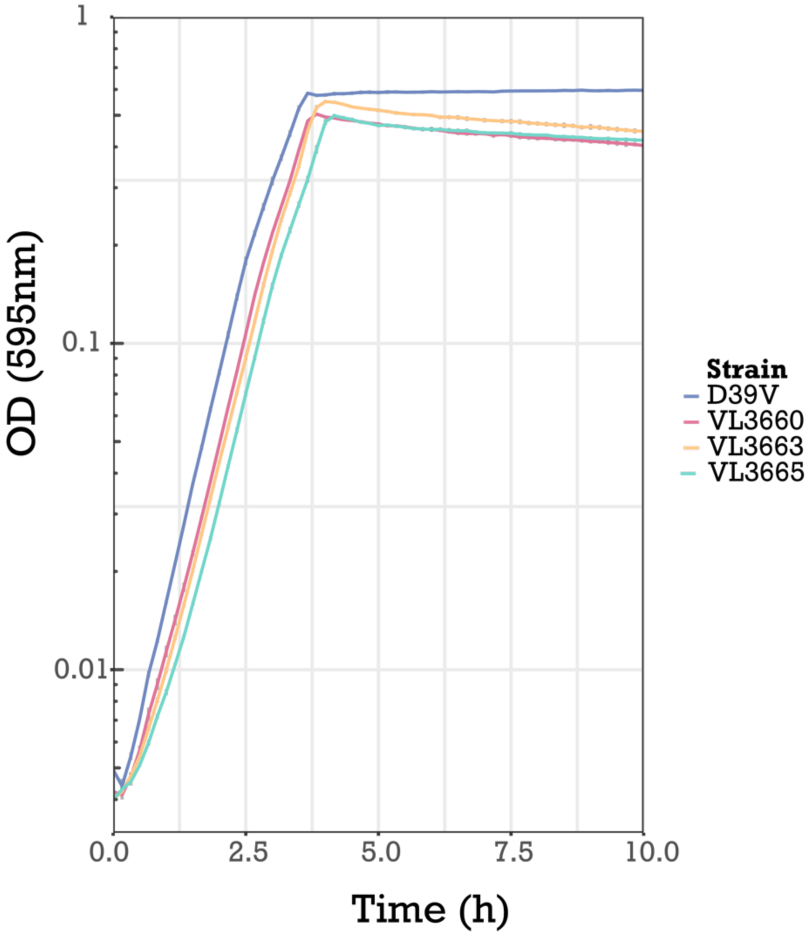


Figure S1: Cell density (OD595) of the bacterial cultures were measured every 10 min. A typical experimental outcome is shown (repeated at least 3 times). The values represent averages of three replicates. D39V = wild type, VL3660 = *Δcps*, VL3663 = *Δcps, Δply,* VL3665 = *Δcps, Δply, ΔlytA*
